# TSPO-PET Reveals Higher Inflammation in White Matter Disrupted by Paramagnetic Rim Lesions in Multiple Sclerosis

**DOI:** 10.1101/2025.01.03.627857

**Authors:** Ceren Tozlu, Keith Jamison, Yeona Kang, Sandra Hurtado Rua, Ulrike W. Kaunzner, Thanh Nguyen, Amy Kuceyeski, Susan A. Gauthier

**Author notes:** **Corresponding Author:** Susan A. Gauthier, DO, MPH **Mailing address:** Judith Jaffe Multiple Sclerosis Center, 1305 York Avenue, New York, NY10021, USA.; **Email:**. Authors contributed equally to this work (Amy Kuceyeski, Susan A. Gauthier).

## Abstract

**Objective:** To explore whether the inflammatory activity is higher in white matter (WM) tracts disrupted by paramagnetic rim lesions (PRLs) and if inflammation in PRL-disrupted WM tracts is associated with disability in people with multiple sclerosis (MS).

**Methods:** Forty-four MS patients and 16 healthy controls were included. 18 kDa-translocator protein positron emission tomography (TSPO-PET) with the ^11^C-PK11195 radioligand was used to measure the neuroinflammatory activity. The Network Modification Tool was used to identify WM tracts disrupted by PRLs and non-PRLs that were delineated on MRI. The Expanded Disability Status Scale was used to measure disability.

**Results:** MS patients had higher inflammatory activity in whole brain WM compared to healthy controls (p=0.001). Compared to patients without PRLs, patients with PRLs exhibited higher levels of inflammatory activity in the WM tracts disrupted by any type of lesions (p=0.02) or PRLs (p=0.004). In patients with at least one PRL, inflammatory activity was higher in WM tracts highly disrupted by PRLs compared to WM tracts highly disrupted by non-PRLs (p=0.009). Elevated inflammatory activity in highly disrupted WM tracts was associated with increased disability in patients with PRL (p=0.03), but not in patients without PRL (p=0.2).

**Interpretation:** This study suggests that patients with PRLs may exhibit more diffuse WM inflammation in addition to higher inflammation along WM tracts disrupted by PRLs compared to non-PRLs, which could contribute to larger lesion volumes and faster disability progression. Imaging PRLs may serve to identify patients with both focal and diffuse inflammation, guiding therapeutic interventions aimed at reducing inflammation and preventing progressive disability in MS.

## Introduction

Understanding the factors driving disease progression in multiple sclerosis (MS) is essential for achieving accurate prognoses and developing new therapeutic targets. Compartmentalized inflammation within the central nervous system (CNS) of patients with MS is believed to play a significant role in advancing the understanding of the disease. Persistent smoldering inflammation in chronic active lesions (CALs) has been linked to progressive ambulatory decline in these patients^1^. Paramagnetic rim lesions (PRLs)—a subset of CALs defined by a dense rim of iron-laden, pro-inflammatory immune cells—can be visualized using gradient echo MRI (GRE). Recent studies have associated PRLs with a more aggressive cognitive and ambulatory impairment^2–6;^ however, it remains uncertain whether this correlation is due to the presence of PRLs themselves or other underlying, unmeasured factors. Patients with PRLs typically show a higher lesion burden^1^, raising the question of whether they experience more diffuse inflammatory activity, which could contribute to greater white matter (WM) damage and a predisposition to form PRLs.

Positron Emission Tomography (PET) in combination with radioligand ^11^C-PK11195 that binds to the 18-kDa translocator protein (TSPO) is a frequently used imaging technique to measure innate immune cell activation such as microglia and macrophages *in vivo*^7^. TSPO PET has provided insights into the mechanisms underlying tissue damage that may not be detectable with MRI, revealing associations between these mechanisms and disability^8,9^. ^11^C-PK11195 PET (PKPET) has been used to demonstrate greater inflammatory activity within PRLs compared to non-PRLs^10^. Recent studies suggest a broader inflammatory phenotype in patients with PRLs, including elevated levels of CSF chitinase 3-like-1, a marker of astrocyte and macrophage/microglial activation^11^. PKPET has provided an opportunity to explore diffuse neuroinflammatory activity throughout the brains of patients with MS^12^; and of particular interest would be to examine inflammation in the normal-appearing WM (NAWM) along the WM tracts adjacent to these pathological lesions.

The Network Modification (NeMo) tool^13^ is an effective approach that uses a large normative tractography dataset to quantify disruption in WM tracts due to brain lesions and was previously used in stroke^14^, traumatic brain injury^15^, and MS ^4,16–18^. We have used the NeMo tool to show that disability in MS patients was more associated with disruptions in WM tracts due to PRLs than disruptions in WM tracts due to non-PRLs^3^. Building on these findings, we hypothesize that the impact of PRLs on disability is partially due to increased inflammatory activity along the WM tracts adjacent to these lesions. Therefore, in this study, we propose a multi-modal approach with PKPET to test for increased neuroinflammatory activity along WM tracts disrupted by any kind of lesion in patients with and without PRLs. Additionally, we aim to assess whether neuroinflammatory activity differs in WM tracts disrupted by PRLs versus non-PRLs within PRL-positive patients only. Finally, we explore the relationship between disability and the inflammatory activity in the disrupted WM tracts in patients with and without PRLs.

## 2. Materials and Methods

### Cohort

This is a cross-sectional retrospective study including 44 patients (age 45.45 ± 13.82, 61.4% female) with a diagnosis of MS, including 27 relapsing-remitting (RR), 6 primary progressive (PP) and 11 secondary progressive (SP). Sixteen healthy controls (HC) (age: 56.19 ± 10.05, 25 % females) were scanned with PKPET for comparison. MRI scans were performed within 15 ± 25 days of the PKPET scans. Inclusion in this analysis required individuals to have both a PKPET scan and an MRI protocol that included quantitative susceptibility mapping (QSM) for PRL detection. Patient characteristics and clinical data were obtained within 1 month of the individual’s brain MRI and PKPET scan. The following information was collected for each MS patient: demographics (age and sex), clinical phenotype (RR, SP, PP), disease duration, Expanded Disability Status Scale (EDSS), and treatment efficacy level. EDSS was used to quantify disability in MS patients. Treatment level was either low efficacy (glatiramer acetate, low and high dose interferon beta -1a, dimethyl fumarate), high efficacy (fingolimod, natalizumab, and ocrelizumab), or none (no treatment or monthly steroid). All studies were approved by an ethical standards committee on human experimentation and written informed consent was obtained from all subjects.

### Image acquisition, processing, and structural connectome extraction

MRI was performed at 3 T using a GE Signa HDxt (50% patients) or Siemens Magnetom Skyra scanner. The MRI protocol consisted of sagittal 3-dimensional (3D) T1-weighted (T1w) sequence for anatomic structure, 2-dimensional (2D) T2-weighted (T2w) fast spin-echo, and 3D T2w fluid-attenuated inversion recovery (FLAIR) sequences for lesion detection, gadolinium-enhanced 3D T1w sequence for acute lesion identification, and axial 3D multi-echo GRE sequence for QSM. The detailed scanning protocols are provided in Zinger et al.^19^ and are overall very similar, including the axial 3D multi-echo GRE sequence for QSM. The harmonized QSM imaging protocol has been demonstrated to have high reproducibility across different scanner vendors^20^. QSM was reconstructed from complex GRE images using a fully automated Morphology Enabled Dipole Inversion algorithm zero-referenced to the cerebrospinal fluid (MEDI+0)^21^.

### PK11195 radioligand production and PET imaging

The radioligand [N-methyl-11C(R)-1-(2-chlorophenyl)-N-(1methylpropyl)-3- isoquinolinecarboxamide], known as ^11^C-PK11195 was prepared by modifying previously described procedures. Briefly, 1 mg (2.95 mol) desmethyl-PK11195 in 350 ml DMSO was treated with 10 mmol aqueous sodium hydroxide and allowed to react with (C-11) methyl iodide. Following the reaction ^11^C-PK11195 was purified by high-performance liquid chromatography and formulated in saline with 7% ethanol. Following intravenous administration of 370–555 MBq (1015 mCi) of 11C-PK11195, dynamic PET scans over a period of 60 min were acquired in list mode with a whole-body PET/ CT scanner (mCT, Siemens/CTI). The PET camera has a spatial resolution of ∼4 mm measured as the reconstructed full width at half-maximum of a point source in air. PET scans were corrected for photon absorption and scatter, using an in-line CT scanner set at 120 kV, a pitch of 1.5, and 30 mA. PET data were reconstructed in a 400 x 400 matrix with a voxel size of 1.082 x 1.082 x 2.025 mm^3^ using a zoom of 2.0 and an iterative + time of flight (TOF) list-mode reconstruction algorithm provided by the manufacturer.

### PET data quantification

PET images were reconstructed into 22 frames (four frames of 15 s each, then 4 x 30 s, 3 x 60 s, 2 x 120 s, 8 x 300 s, and 1 x 600 s). PET images were co-registered with their corresponding MRI scans using PMOD^®^ (PMOD Technologies Ltd). Innate immune cell activity expressing TSPO was assessed by measuring the specific binding of ^11^C -PK11195, using the distribution volume ratio (DVR) across the entire brain (DVR map)^22^. To estimate the DVR PKPET map, the Logan reference method was employed, utilizing the radioactivity concentrations in the brain and a reference region. The reference region, essential for calculating DVR, was defined using the super clustering method^23^ which was conducted using PMOD.

DVR PKPET images were transformed to the individual’s T1 native space using the inverse of the T1 to GRE transform and trilinear interpolation. Individual T1 images were then normalized to MNI space using FSL’s linear (FLIRT) and nonlinear (FNIRT) transformation tools (http://www.fmrib.ox.ac.uk/fsl/index.html); transformations with trilinear interpolation were then applied to transform both native anatomical space DVR PKPET images to MNI space. The transformations were concatenated to minimize interpolation. DVR PET images were visually inspected after the transformation to MNI space to verify accuracy. The average DVR from PKPET across the WM in HC and MS patients were presented in Supplementary Figure 1.

### Lesion segmentation and PRL identification

Lesion masks were produced for each MS patient in a semi-automated process. The WM hyperintensity lesion masks were created by running the T2 FLAIR images through the Lesion Segmentation Tool (LST) and were further hand-edited as needed. T2 FLAIR-based lesion masks were transformed to the individual’s T1 native space using the inverse of the T1 to GRE transform and nearest-neighbor interpolation. Individual T1 images were then normalized to MNI space using FLIRT and FNIRT transformation tools; transformations with nearest-neighbor interpolation were then applied to transform the native anatomical space T2FLAIR lesion masks to MNI space. The transformations were concatenated (T2FLAIR to T1 to MNI) to minimize interpolation effects. Lesions were visually inspected after the transformation to MNI space to verify the accuracy of coregistration and lesion volume (in mm^3^) calculated. The NAWM mask was created by subtracting the lesion mask from the WM mask.

The consensus of two blinded reviewers was used to identify PRLs on QSM^24^, and a third independent reviewer resolved any discrepant lesions. Lesions with partial or complete rims were considered PRLs, in accordance with the most recent consensus statement on imaging chronic active lesions^25^.

### Network Modification Tool

The MNI space non-PRL and PRL masks were processed through the newest version of the NeMo Tool^13^, which provides voxel-level estimates of how much the lesions disrupt streamlines passing through WM voxels or connecting to GM voxels. The NeMo Tool calculates a voxel’s disruption from a lesion mask as the percent of streamlines in a tractography reference set connecting to or passing through that voxel that also passes through the lesion mask. The tractography reference set is based on diffusion MRI data from 420 unrelated healthy controls (206 female, 214 male, 28.7 ± 3.7 years). The disruption in the WM tracts ranges between 0 and 1, where 0 represents no disruption and 1 represents complete disruption. WM voxel disruptions were calculated separately for T2 FLAIR and non-PRL masks for all patients (N=44) and PRL masks for only those with at least one PRL lesion (N=26) (Supplementary Figure 2 for the lesion and WM disruption masks).

### White matter masks

The WM mask was created using the HCs only to standardize the WM mask across all participants and minimize variability due to MS-related WM alterations. The WM masks were obtained from the aseg files extracted from FreeSurfer in 16 HCs. The WM masks were then coregistered to the MNI space using the transformation matrix which was used to coregister the T1 images to the MNI space (see above for more details). The final WM mask template was created using the voxels that existed in more than 50% of the HCs (See Supplementary Figure 3 for the probability WM mask and the thresholded version). The DVR PKPET and WM disruption metrics were then overlapped to the WM mask and measured on the WM mask after removing the WM voxels within the 1mm proximity of ventricles and gray matter.

### Statistical Analysis

Based on the normality of the data as assessed with the Shapiro-Wilk test, the Student’s t-test or Wilcoxon rank sum test was used to compare the demographics or neuroimaging metrics (Table 1). The unpaired student’s t-test was used to compare the voxel-wise DVR metrics between healthy controls and MS patients as well as between patients with vs without PRL (see Figure 1, panels A and C). Additionally, we also compared the voxel-wise DVR between the groups using ANCOVA where age and sex were used as a covariate (See Supplementary Figure 4). Next, z-scores for DVR metrics were calculated for each voxel using 16 healthy controls as reference (Figure 1, Panel B). Subject-level averaged DVR metrics were compared between HC, all MS patients, MS patients with PRL, and MS patients without PRL (Figure 1, Panel D) using ANCOVA where age and sex were used as covariates.

**Figure 1:**
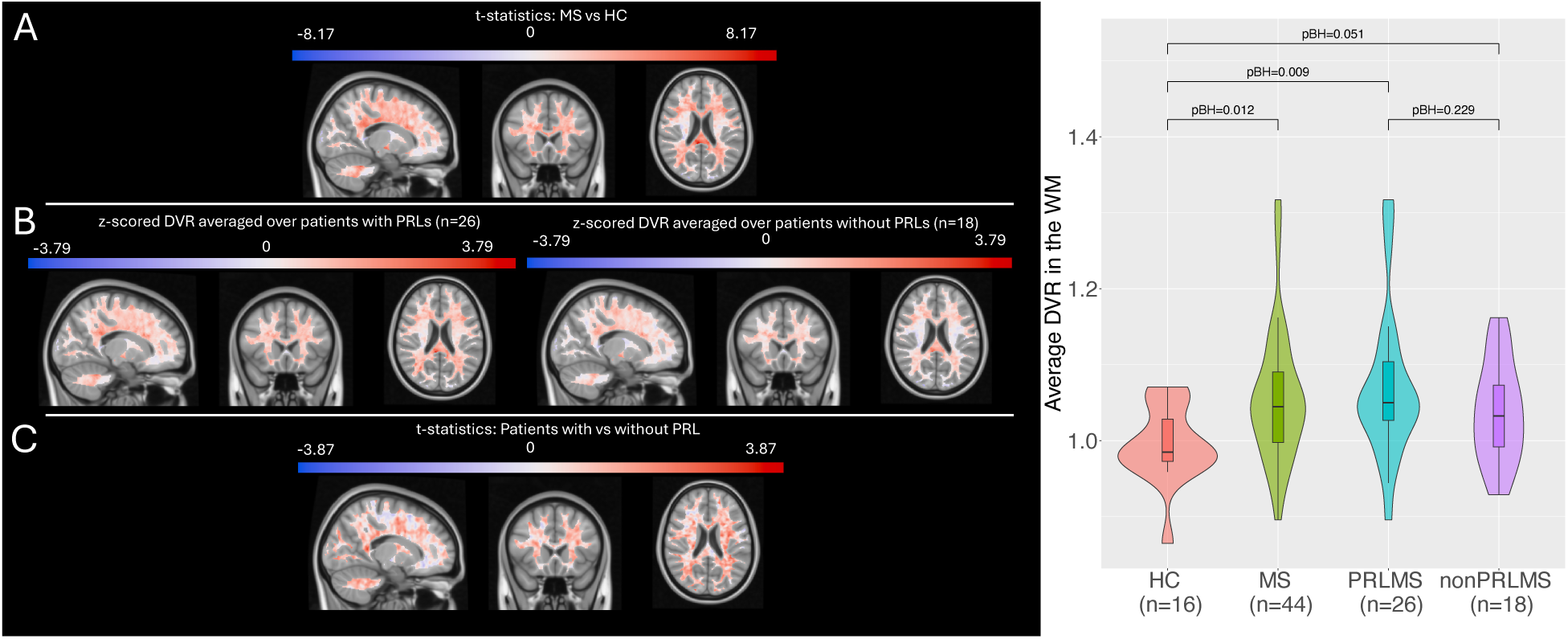
(A) Statistics comparing voxel-wise DVR between MS patients and healthy controls. The statistics values were derived from the Student’s t-test analysis, which compared voxel-wise DVR between MS patients and healthy controls. Positive statistics values indicate higher DVR in MS patients compared to healthy controls. (B) The z-scored voxel-wise DVR metrics in patients with and without PRL, separately. The z-scoring was obtained using the voxel-wise DVR metrics in 16 healthy controls. (C) Statistics comparing voxel-wise DVR between MS patients with vs without PRL. The statistics values were derived from the Student’s t-test analysis, which compared voxel-wise DVR between MS patients with vs without PRL. Positive values indicate higher DVR in MS patients with PRL compared to those without PRL. The voxels within 1mm proximity to CSF and cortex were excluded. (D) The average DVR metrics at the subject level for HC, all MS patients, MS patients with PRL, and MS patients without PRL. ANCOVA was used to compare the average DVR between MS patients and healthy controls as well as between the MS patients with vs without PRL. Age and sex were used as a covariate in the ANCOVA models. The BH-adjusted p-values (pBH) were presented in the figures.

**Table 1:**
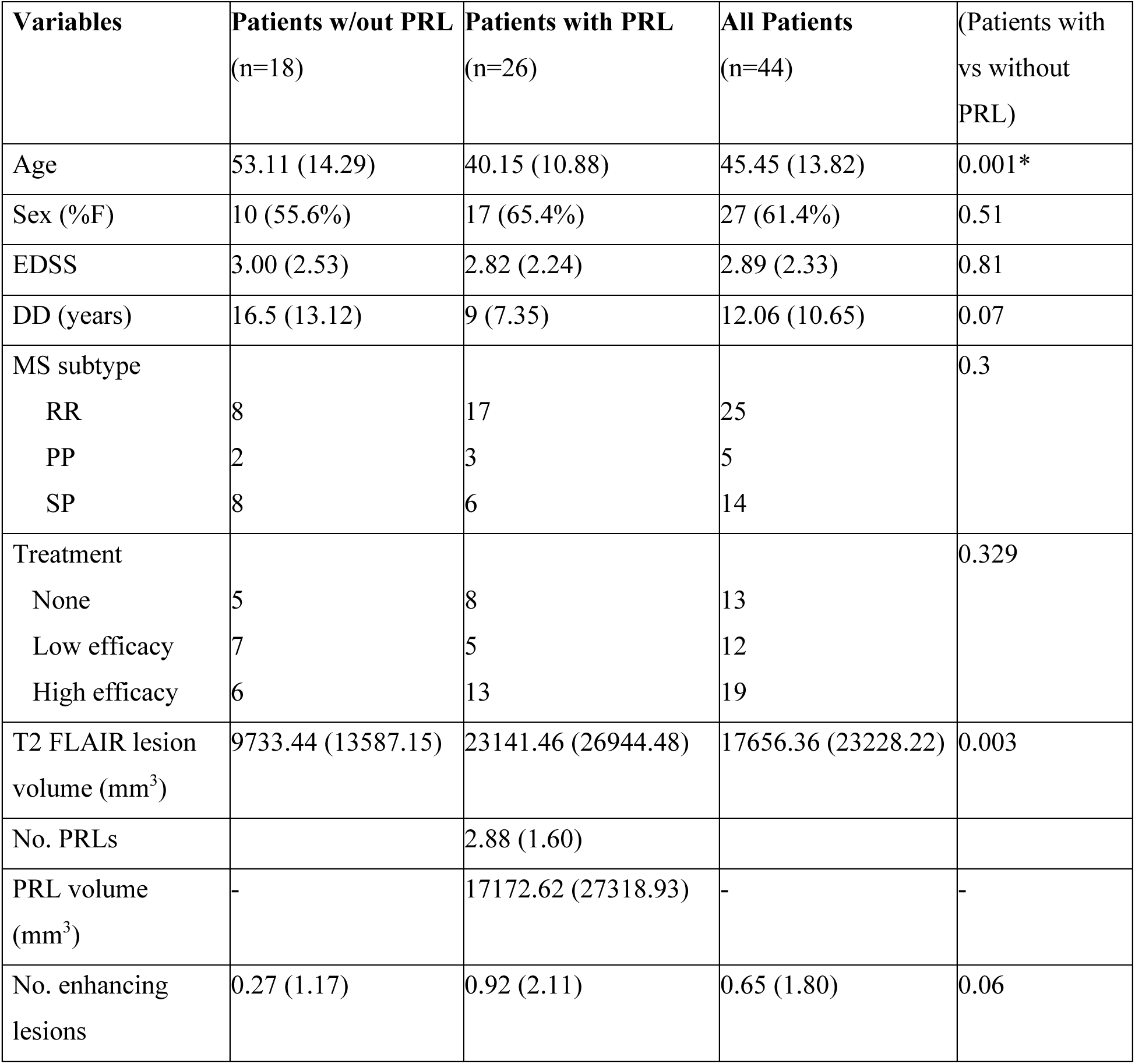
Clinical and MRI Characteristics of all Patients. Values were presented as mean (standard deviation). Based on the normality of the data, the Student’s t-test or Wilcoxon rank sum test was used to test the group differences. *indicates significance at p < 0.05. Abbreviations: DD, disease duration; RR, relapsing-remitting; SP, secondary progressive; PP, primary progressive; EDSS, expanded disability status scale; FLAIR, fluid-attenuated inversion recovery; PRL, paramagnetic rim lesion. The treatment information was categorized as low efficacy (glatiramer acetate, low and high dose interferon beta -1a, dimethyl fumarate), high efficacy (fingolimod, natalizumab, and ocrelizumab) or no treatment (no treatment or monthly steroid) groups.

The average of the z-scored DVR metrics was calculated in the WM tracts with varying levels of lesion-based disruption for each subject (See Figure 3). A voxel’s disruption level was either no disruption (score of 0), low disruption (scores between 0 and 0.1), or high disruption (scores between 0.9 and 1). For the patients with PRLs, the average of the z-scored DVR metrics was calculated for WM tracts with varying levels of disruption due to PRLs and non-PRLs masks separately. ANCOVA was used to compare the average of z-scored DVR in the WM tracts with varying levels of disruption between the patients with vs without PRL. Age, sex, EDSS, and MS type were used as covariates in this ANCOVA analysis. A paired t-test was used to compare the average of z-scored DVR in the WM tracts with varying levels of disruption due to PRLs vs non-PRLs in patients with PRL. Multiple comparison p-value correction was performed using Benjamini-Hochberg^26^.

**Figure 2:**
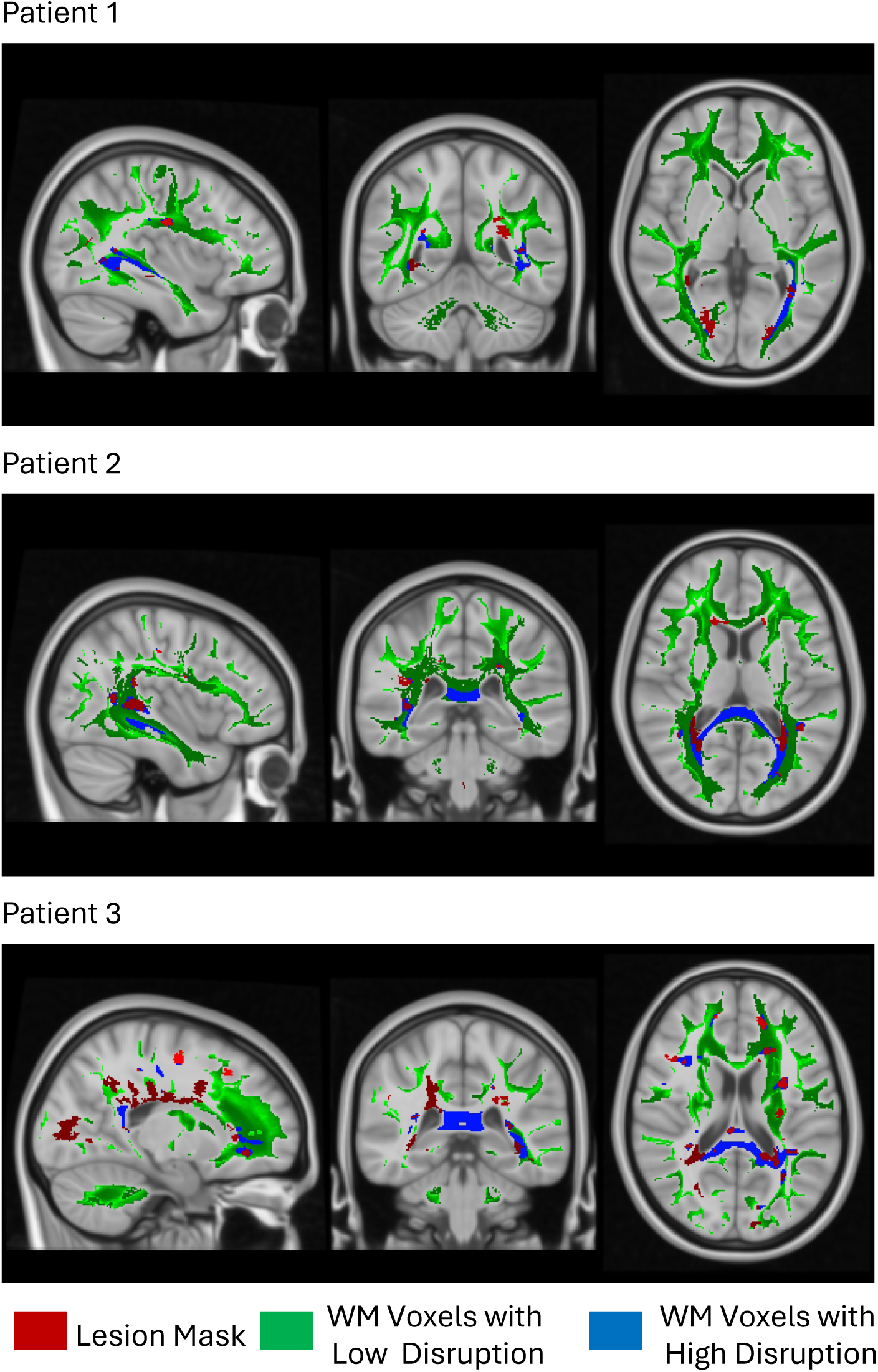
Examples from three different MS patients representing the lesion mask (red), white matter voxels with low disruption (green) due to any type of WM lesion, and white matter voxels with high disruption (blue) due to any type of WM lesion.

**Figure 3:**
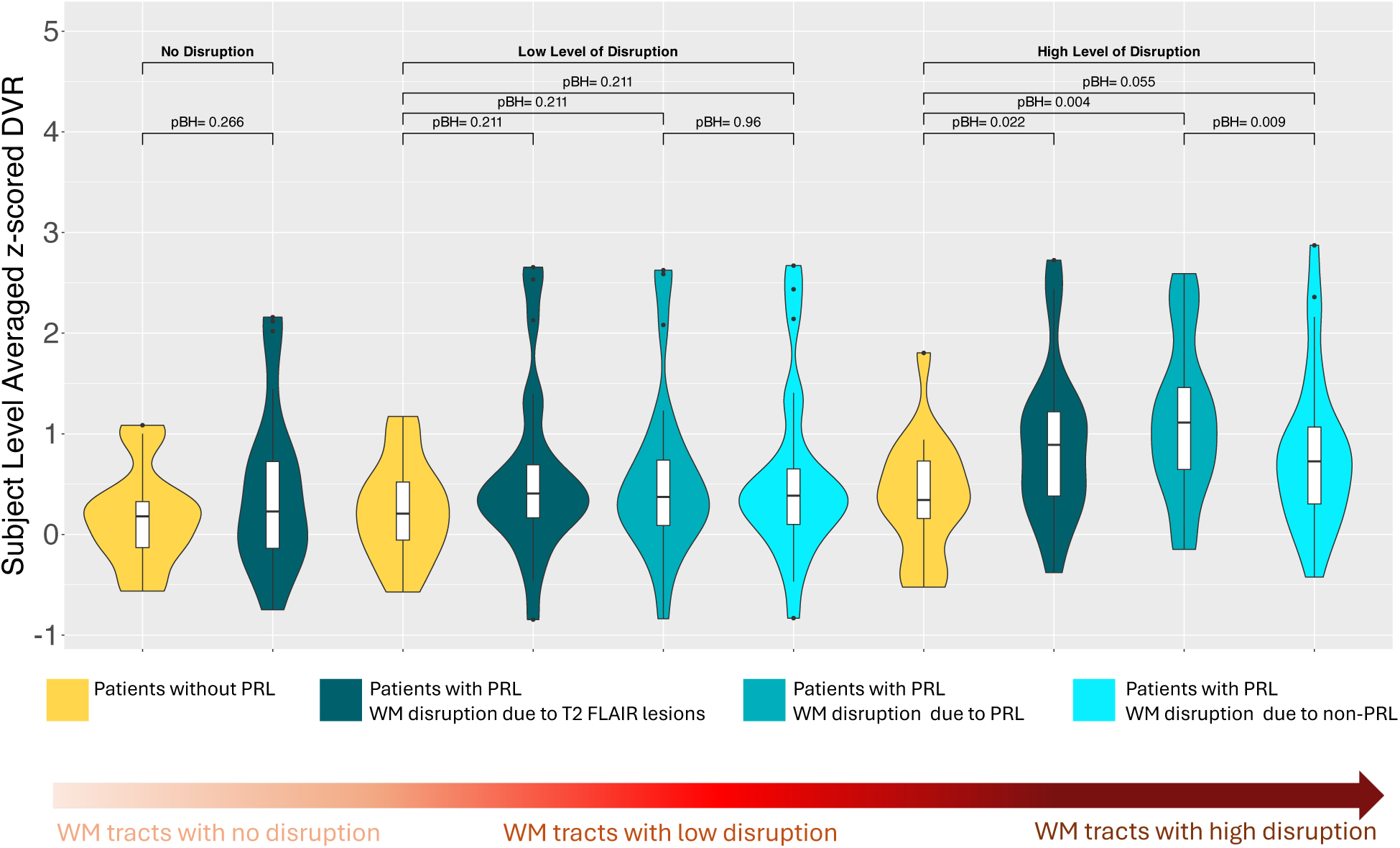
The patient-level z-scored DVR metrics averaged across the WM tracts where i) there was no disruption due to MS lesions (i.e. disruption in the WM tracts is zero), ii) there was a low level of disconnection due to lesions (i.e. disruption in the WM tracts is between 0 and 0.1), and iii) there was a high level of disconnection due to lesions (i.e. disruption in the WM tracts is between 0.9 and 1). The voxel-wise z-scored DVR was created in the WM using the voxel-wise DVR in 16 healthy controls. Then, the patient-level averaged z-scored metrics were calculated for each MS patient separately. The disruption metrics were separated as disruption due to PRL and disruption due to non-PRL for patients with at least one PRL. The yellow color represents the patients without PRL and the blue colors represent the patients with at least one PRL. The comparisons between the patients with vs without PRL were performed with ANCOVA where age, sex, and MS type were controlled. In patients with PRL, a paired t-test was used to compare the average of z-scored DVR in the WM tracts with varying levels of disruption due to PRLs vs non-PRLs. The BH-adjusted p-values (pBH) were presented in the figures.

A linear model was used to identify the association between the EDSS and the averaged z-scored DVR metrics in the disrupted WM tracts at varying levels in patients with and without PRLs separately as well as in all MS patients (See Figure 4). Age, sex, and MS type were used as covariates in these linear models. The p-values derived from these linear models were corrected using the Benjamini-Hochberg method^26^.

**Figure 4:**
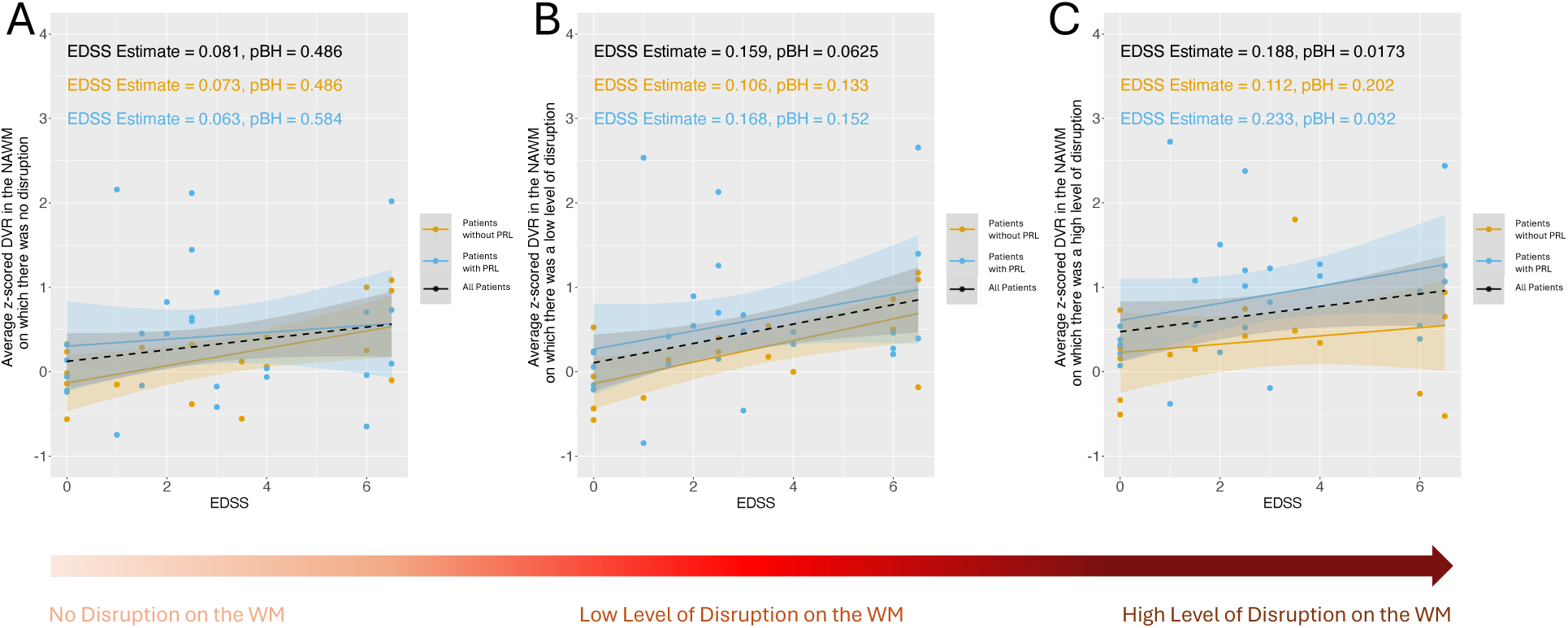
The scatterplots of EDSS and subject-level averaged z-scored DVR for patients without PRL (orange color), patients with PRL (blue color), and all patients (black color). We computed the subject-level averaged z-scored DVR in the WM tracts where A) there was no disruption due to MS lesions (i.e. disruption in the WM tracts is zero), B) there was a low level of disruption due to MS lesions (i.e. disruption in the WM tracts is between 0 and 0.1), and C) there was a high level of disruption due to MS lesions (i.e. disruption in the WM tracts is between 0.9 and 1). The estimates and the p-values associated with EDSS were calculated using a linear model where the output was the subject-level averaged z-scored DVR and the inputs were EDSS, age, sex, and MS type. The BH-adjusted p-values (pBH) were presented in the figures.

The complete dataset of HCs was not age- or sex-matched. To show the consistency of our findings, we created age and sex-matched HCs (age: 53.36 ± 9.7; 4 females and 7 males) and z-scored the DVR metrics using these 11 HCs. The *matchit* function from the MatchIt package in R was used with nearest neighbor matching and mahalanobis distance to create 11 HCs that are age- (Wilcoxon Rank Sum test p-value=0.66) and sex- (chi-square test p-value=0.199) matched to the MS cohort. The replicated findings, which are very similar to the main results using all 16 healthy controls, were presented in the Supplementary Information. We chose to present the results based on the larger healthy control cohort in the main text, so we had a larger sample size to enhance statistical power and variability.

## 3. Results

### 3.1. Patient Characteristics

Table 1 presents the clinical and neuroimaging characteristics of all 44 patients with MS. Although there were no differences in disability, patients with PRLs were significantly younger (p = 0.001) and had a higher T2 FLAIR lesion volume (p = 0.001) compared to those without PRLs. There were no significant differences in non-PRLs volume or gadolinium-enhancing lesion count between patients with and without PRLs.

### 3.2. TSPO binding within the WM across patients and controls

MS patients exhibited higher DVR values in most WM voxels compared to HC, particularly in the periventricular regions (Figure 1, A). Voxel-wise analysis revealed that both MS subgroups— those with and without PRLs—had higher DVR values than HC (Figure 1, B), and that PRL-patients demonstrated relatively greater DVR than PRL-negative patients (Figure 1, C). At the subject level, average DVR across the WM was significantly higher in all MS patients compared to HC, and this pattern held true for both patient subgroups (Figure 1, D). However, the difference in DVR between PRL-positive and PRL-negative patients was not statistically significant at the subject level.

### 3.3. TSPO binding in the WM tracts with varying levels of lesion disruption

The NeMo tool was used to measure MS lesion-based WM voxel disruption, which is defined as the percentage of WM streamlines passing through that voxel and also intersect the lesion mask. Figure 2 illustrates the WM voxels with low and high disruption by MS lesions in three different MS patients. As expected, the percentage of WM voxels with low disruption was greater than those with high disruption. Highly disrupted WM voxels were observed both near and distant from the lesions, suggesting that voxels distant from lesions can still exhibit high disruption if they are strongly connected to the lesions.

For the patients with PRLs, voxel-level WM disruption was computed for all MS lesions as well as for PRLs and non-PRLs masks separately. There were no significant differences observed between patients with vs without PRLs in the z-scored DVR for WM tracts with no disruption (p=0.26) or low disruption (adjusted p=0.21) due to MS lesions (Figure 3). However, WM tracts with high disruption in patients with PRLs showed higher z-scored DVR compared to WM tracts with high disruption in patients without PRLs (adjusted p=0.022). Additionally, among patients with PRLs, z-scored DVR was higher in WM tracts disrupted by PRL compared to WM tracts disrupted by non-PRLs (adjusted p= 0.009). The results were consistent when age and sex-matched HCs were used to z-score the DVR metrics in MS patients (See Supplementary Figure 5). Patients with PRLs showed higher z-scored DVR in highly disrupted WM tracts compared to highly disrupted WM tracts in patients without PRLs (adjusted p=0.028). Among patients with PRLs, the z-scored DVR was higher in WM tracts disrupted by PRL compared to WM tracts disrupted by non-PRLs (adjusted p= 0.015).

### 3.4. The relationship of disability and TSPO binding in WM tracts

Finally, we investigated the relationship between EDSS and DVR in patients with and without PRLs. The average of z-scored DVR was computed on the WM tracts with no disruption (Figure 4, A), low disruption (Figure 4, B), and high disruption (Figure 4, C), separately. No significant association was found between EDSS and z-scored DVR in WM tracts with no or low disruption in either MS group or in the combined MS group. However, there was a significant association between EDSS and z-scored DVR in highly disrupted WM tracts across all MS patients (adjusted p=0.017) and in patients with PRLs (adjusted p=0.0320). There was no significant association for patients without PRLs (adjusted p=0.202). When the age and sex-matched HCs were used to z-score DVR metrics in MS patients, we found similar results (See Supplementary Figure 6). That is, there was a significant association between EDSS and z-scored DVR in highly disrupted WM tracts across all MS patients (adjusted p=0.0147) and within patients with PRLs (adjusted p=0.0329), but not in patients without PRLs (adjusted p=0.194).

## 4. Discussion

This study used a novel multi-modal PET and MRI approach to investigate patterns of neuroinflammatory activity in the WM of MS patients. We demonstrated that neuroinflammatory activity was elevated in WM tracts highly disrupted by any kind of lesion in patients with PRLs compared to patients without. In patients with PRLs, we found that WM highly disrupted by PRL lesions had higher inflammation than WM highly disrupted by non-PRL lesions. Furthermore, increased inflammation was associated with greater disability in patients with PRLs. These findings provide further evidence that PRLs may be linked to a distinct pattern of neuroinflammatory activity, which could enhance our understanding of its association with higher disability levels in MS.

Previous studies that utilized TSPO PET to measure neuroinflammatory activity in the NAWM of MS patients compared to HCs have yielded contradictory results. While some studies reported higher neuroinflammatory activity in NAWM MS patients with RR and progressive MS^27–29^ or in clinically isolated syndrome (CIS)^9^, other studies found no difference in neuroinflammatory activity in NAWM of RRMS patients^30–32^. Various factors likely contribute to these inconsistent findings, including variations in patient cohorts, TSPO ligands, and PET processing and quantification approaches. In addition, these studies focused on the NAWM as a whole, whereas our study examined specific WM tracts disrupted by lesions as identified using the NeMo Tool.

Previous studies have shown WM disruption in all NAWM as measured by reduced fractional anisotropy (a summary metric of diffusion MRI) is associated with increased NAWM TSPO uptake^30,32^. However, this work analyzed the average fractional anisotropy of all NAWM, whereas our approach offers a more specific analysis by analyzing the level of neuroinflammation across varied levels of WM disruption. Although one could use the spatial proximity of WM voxels to lesions or CSF^33^, it’s important to note two considerations: WM voxels close in proximity to a lesion may not share streamlines with said lesion (i.e. may be on crossing or kissing WM tracts) and WM voxels disrupted by a lesion may not be in close proximity to that lesions (See Figure 2). Therefore, it is important to consider the topology of the lesions and how they disrupt WM voxels of interest.

We demonstrate that MS patients with at least one PRL exhibit higher inflammatory activity in highly disrupted WM tracts compared to those without PRLs. To validate the hypothesis that PRLs might contribute to the inflammatory activity along the WM tracts that connect them to other brain regions, we also compared the inflammatory activity on the WM tracts impacted due to PRLs vs non-PRLs among the patients with PRL only. Consistent with our results indicated above, higher inflammation was observed in the WM tracts highly disrupted due to PRLs compared to those without PRLs. These findings suggest that even within the same patient, the neuroinflammatory activity in the WM tracts differs based on the presence of PRL. These findings expand our previous work, which demonstrated that disability in MS patients was more accurately predicted by disruptions in WM tracts due to PRLs than by those caused by non-PRLs. Together, these results may further explain the higher lesion burden, brain volume loss as well as greater levels of both cognitive and motor disability observed in these patients^1–6,34^

The pathological rim in a PRL evolves for years^35,36^ and provides a substrate for ongoing demyelination and axonal damage.^36,37^ PRLs are notably destructive, with a tendency to slowly expand over time^38^. Considering the pathology of PRLs, there are two possible mechanisms to explain our findings. Patients with PRLs may exhibit heightened global inflammatory activity, as suggested by our current and several previous findings in PRL positive patients, including enlarged choroid plexus^39^, which is considered an immune reserve, elevated CSF chitinase 3-like-1^11^, and leptomeningeal enhancement^40^. Alternatively, damage caused by PRLs might trigger an inflammatory response within WM voxels disrupted by the PRL. This is supported by the increased expression of inflammatory markers in iron-laden, inflamed microglia at the edges of CALs, along with an expanded population of upregulated inflammatory astrocytes^41^ and evidence of increased damage to the periplaque WM^42^. While the exact mechanism remains unclear, our results suggest that the latter hypothesis is more plausible, given that differences between patients with and without PRLs appeared to be driven by PRL-affected WM. Increased inflammatory activity in these highly disrupted WM tracts may impede remyelination, potentially explaining why PRLs tend to expand and ultimately lead to greater overall damage.

Iron may not be present at the rim of all CAL, thus identification of PRLs with GRE MRI may underestimate the true burden of these pathological lesions. Several TSPO PET studies have proposed to capture a broader classification of CALs which includes both rim-active and homogeneously (or whole) active lesions, based on a per-voxel threshold above healthy controls^27,43,44^. Interestingly, these studies consistently report a similar proportion of lesion subtypes, with homogenously (or whole) active lesions reaching nearly half of the total chronic lesion count. Recent work compared this TSPO-based lesion classification with PRLs and found that while PRLs correlated with levels of active lesions, it was the whole-active lesions with the strongest association with disability^44^. While these lesions clearly impact the disease, it is important to note that only PRLs have been extensively validated as having a dense rim of iron-laden, pro-inflammatory immune cells^1,38,45^. In contrast, TSPO-based lesion classification may be confounded by the specificity of TSPO binding, as it is known to bind not only to pro-inflammatory microglia but also to other glial cells, such as astrocytes. This lack of specificity could also influence our results, as TSPO binding may reflect cellular density rather than activation state^46^. Therefore, while TSPO PET offers valuable insights, these findings must be interpreted with caution.

As with most PET studies, a major limitation of this study is the relatively small sample size, which can impact the generalizability of the findings. PET imaging is expensive, time-consuming, and often involves complex logistics related to radiotracer synthesis and patient preparation, limiting PET studies to single-site studies of small cohorts. A larger multi-center study, focusing on a diverse patient population, is needed to validate our findings and improve their generalizability. However, multi-site TSPO PET studies in MS have not yet been published, primarily due to the need for multi-site harmonization. Despite this, there is growing interest in such studies, given the potential need for imaging biomarkers to guide therapeutic interventions targeting diffuse neuroinflammation. Another limitation of our study was that the WM voxels surrounding the MS lesions might have impacted by the partial volume effect, potentially leading to an overestimation of neuroinflammatory activity. However, Figure 2 shows that highly disrupted WM voxels were present both near and far from the lesions, demonstrating that higher inflammatory activity was not confined to areas adjacent to lesions, but was also present in the WM tracts distant from them. A further limitation of this study is that the database used in the NeMo tool consisted of younger, healthy individuals (ages 21 to 35) compared to the MS patients in this study, whose mean age was 45.45 ± 13.82 years. Although the MS patients were older, our previous work showed that the WM tracts estimated via the NeMo can predict disability as well as the WM tracts derived from diffusion MRI^16^.

Future research should aim to longitudinally assess the progression of PRLs and their impact on WM tracts and neuroinflammatory activity. Tracking these changes could provide valuable insights into how the age of PRLs contributes to ongoing neurodegeneration and disease progression in MS, potentially helping to identify critical periods for intervention. Moreover, future studies should explore therapeutic strategies that specifically target the neuroinflammation associated with PRLs. While previous research has yielded mixed results regarding the reduction of signal at the rim of PRLs—whether through quantification of susceptibility changes^19^ or visual reductions on phase imaging^47^—future work could expand to investigate the effectiveness of treatments targeting both the rim of PRLs and the adjacent disrupted WM.

In summary, our study highlights that a more aggressive disease phenotype in patients with MS having PRLs may be linked to heightened inflammatory activity in WM tracts disrupted by these pathologic lesions. These findings suggest that PRLs could serve as a valuable biomarker for identifying patients who might benefit from therapeutic interventions aimed at reducing inflammation and preventing further damage, potentially improving outcomes for individuals with MS. Specifically, PRLs may serve as a practical screening tool for early-stage clinical trials, such as those involving TSPO PET studies, by reducing reliance on PET imaging for prescreening patients with elevated neuroinflammation and improving the feasibility of study designs. Overall, these results contribute to the growing body of literature suggesting that patients with PRLs represent a distinct disease subtype in MS. Through this work, we have deepened our understanding of PRLs, their role in neuroinflammatory patterns and their association with disability in MS.

## Acknowledgments

C.T. performed the coregistration of the lesion masks and PET images to the MNI space, designed the study, performed the statistical analysis, and wrote the manuscript.

K.J. calculated the voxel-wise WM tract disruption maps, helped with the coregistration of the lesion masks and PET images to the MNI space, and edited the article.

Y.K. calculated the DVR metrics from the PET images and edited the article.

S.H.R. helped with the statistical analysis decisions.

U.K. manually edited the lesion masks, collected patient data, and edited the article.

A.K. designed and supervised the study, helped create the NeMo Tool, and edited the article.

S.G. designed and supervised the study, collected the data, and edited the article.

## Data/code availability statement

The deidentified data that support the findings of this study are available upon reasonable request from the corresponding author. The codes to perform the statistical analyses and create the figures are publicly available https://github.com/cerent/PET_NeMo.

## Ethics statement

All studies were approved by an ethical standards committee of Weill Cornell Medicine on human experimentation, and written informed consent was obtained from all patients.

## Supplementary Information

The average DVR from PKPET across the WM was lower in healthy controls (HC) compared to all MS patients. MS patients with PRLs showed a greater increase in DVR than those without PRLs, particularly within the deeper subcortical NAWM.

**Supplementary Figure 1:**
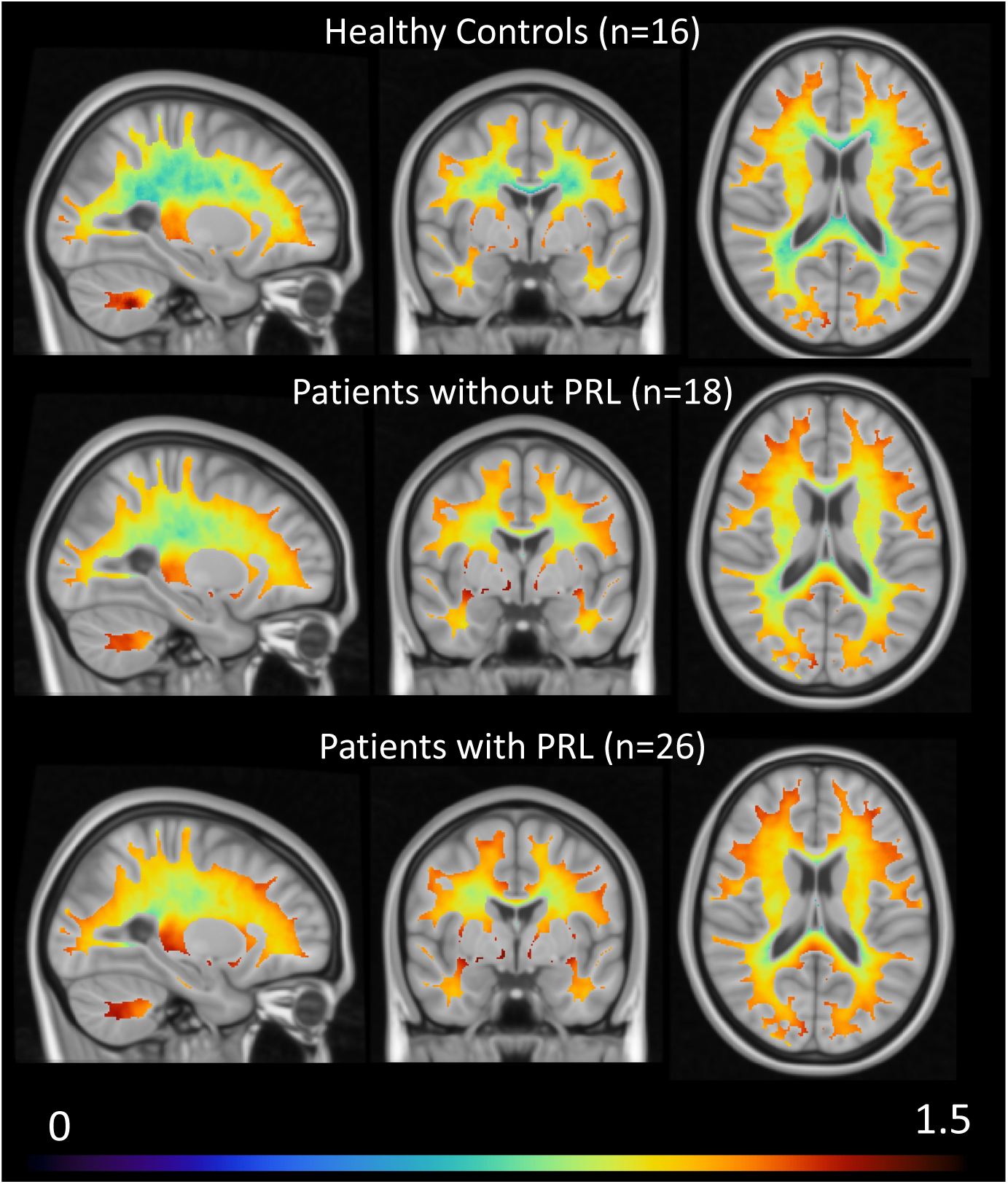
The average DVR maps in the WM in healthy controls, patients with PRL, and patients without PRL. The voxels within 1 mm of proximity from ventricles and gray matter were excluded. The color bar represents the average DVR metrics.

Supplementary Figure 2 represents the T2 FLAIR lesion masks for all patients as well as the PRL and non-PRL lesion masks for the patients with at least one PRL (Panel A). The T2 FLAIR lesion burden was higher in patients with PRL compared to those without PRL. In patients with PRL, the PRL burden was higher compared to the non-PRL burden. Panel B shows that the disruption in the WM tracts due to T2 FLAIR lesions was also higher in patients with PRL compared to those without PRL.

**Supplementary Figure 2:**
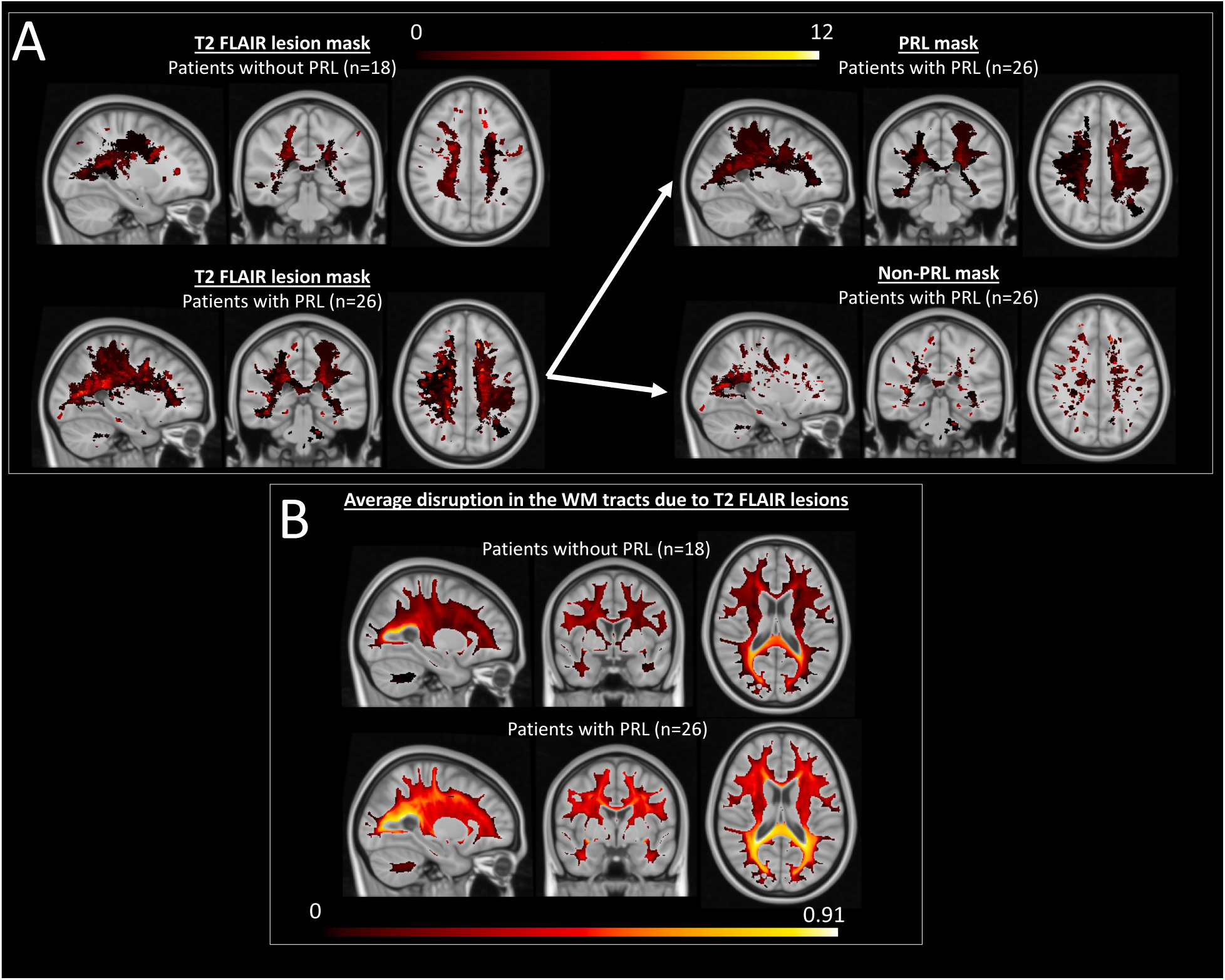
(A) The left panel represents the T2 FLAIR lesion mask in patients with (N=26, figure on the top) and without PRL (N=18, figure on the bottom). The right panel represents the PRL and non-PRL masks in patients with PRL (N=26). (B) The average disruption in the WM tracts due to T2 FLAIR lesions in patients with (N=26, figure on the top) and without PRL (N=18, figure on the bottom). The voxels within 1 mm of proximity from ventricles and GM were excluded from the figures.

Supplementary Figure 3 depicts the process that we used to create the WM mask, which was used then to measure the DVR in the normal-appearing WM. WM mask is created using the voxels that were present in at least 50% of the healthy controls.

**Supplementary Figure 3:**
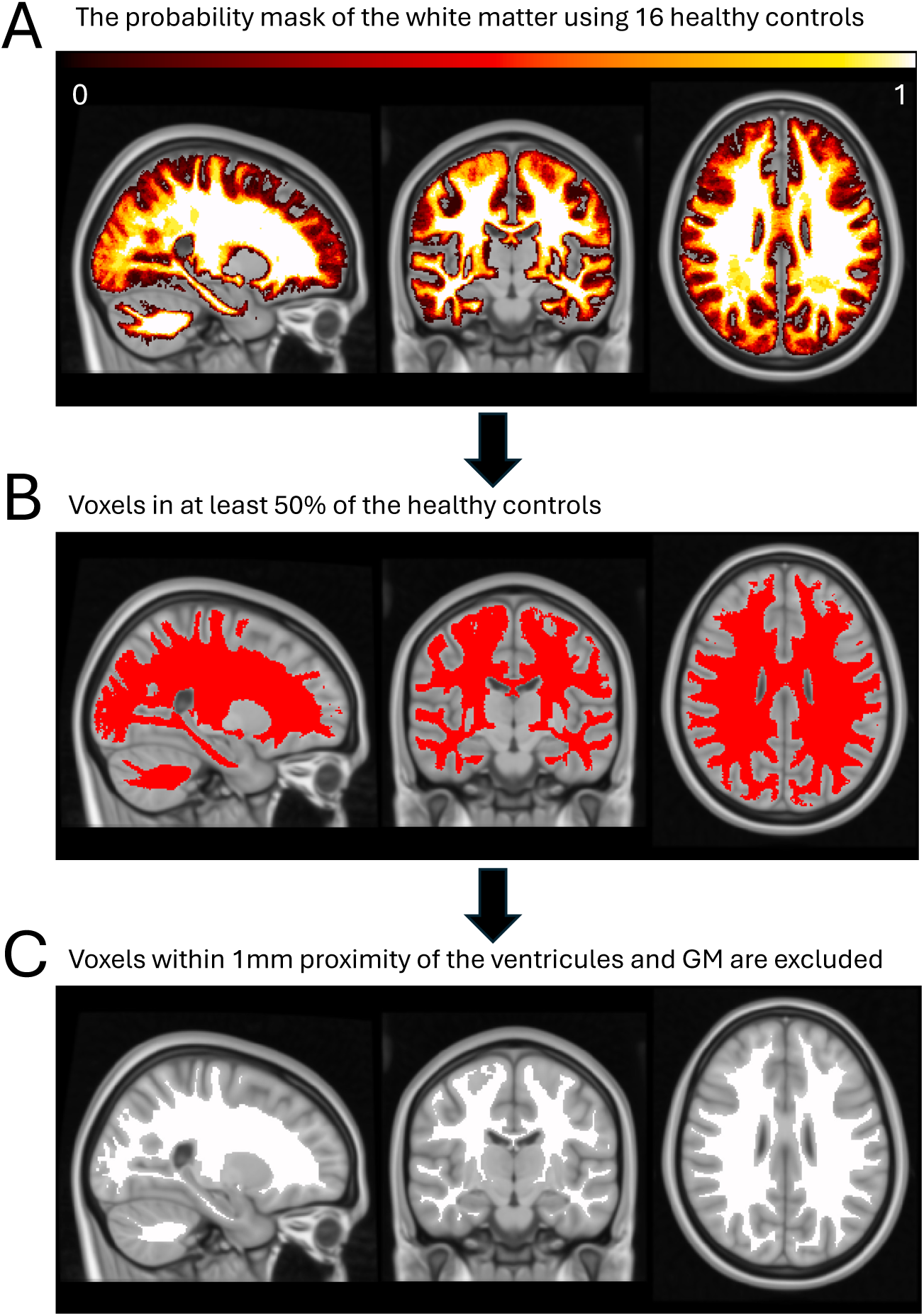
(A) Probability mask generated using the WM segmentations in MNI space from 16 healthy controls. The probability mask shows the percentage of healthy controls with WM segmentation at each voxel, i.e. 0.8 indicates 80% of the healthy controls had WM on that voxel. (B) The thresholded version of the probability mask, including only WM voxels present in at least 50% of the healthy controls. (C) Refined WM mask after excluding voxels within 1mm of the ventricles and gray matter. This final mask was used in our analyses.

The results from Figure 1 - that showed most voxel-wise DVR is higher in MS patients compared to controls as well as in patients with PRL compared to those without PRL - are consistent when ANCOVA is applied using age and sex as covariates.

**Supplementary Figure 4:**
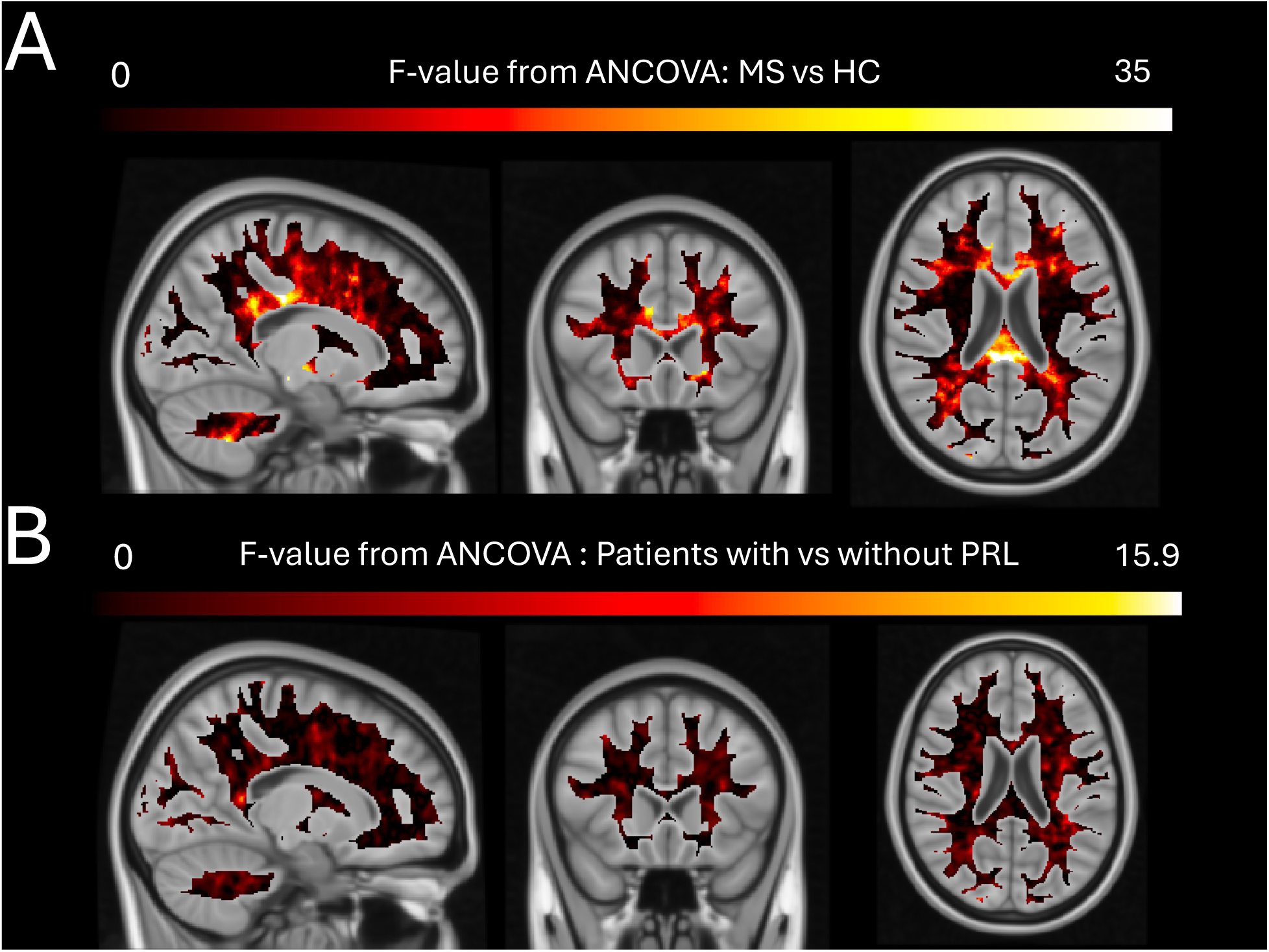
(A) F-value from ANCOVA comparing voxel-wise DVR between MS patients and healthy controls. (B) F-value from ANCOVA comparing voxel-wise DVR between MS patients with vs without PRL. Age and sex were added as covariates in both ANCOVA analyses. The voxels within 1mm proximity to CSF and cortex were excluded.

The results were consistent when age and sex-matched HCs were used to z-score the DVR metrics (See Supplementary Figure 5). Patients with PRLs showed higher z-scored DVR in WM tracts with a high level of disruption due to MS lesions compared to patients without PRLs (adjusted p=0.028). Among patients with PRLs, the z-scored DVR was higher in WM tracts disrupted due to PRL compared to WM tracts disrupted due to non-PRLs (adjusted p= 0.0.15).

**Supplementary Figure 5:**
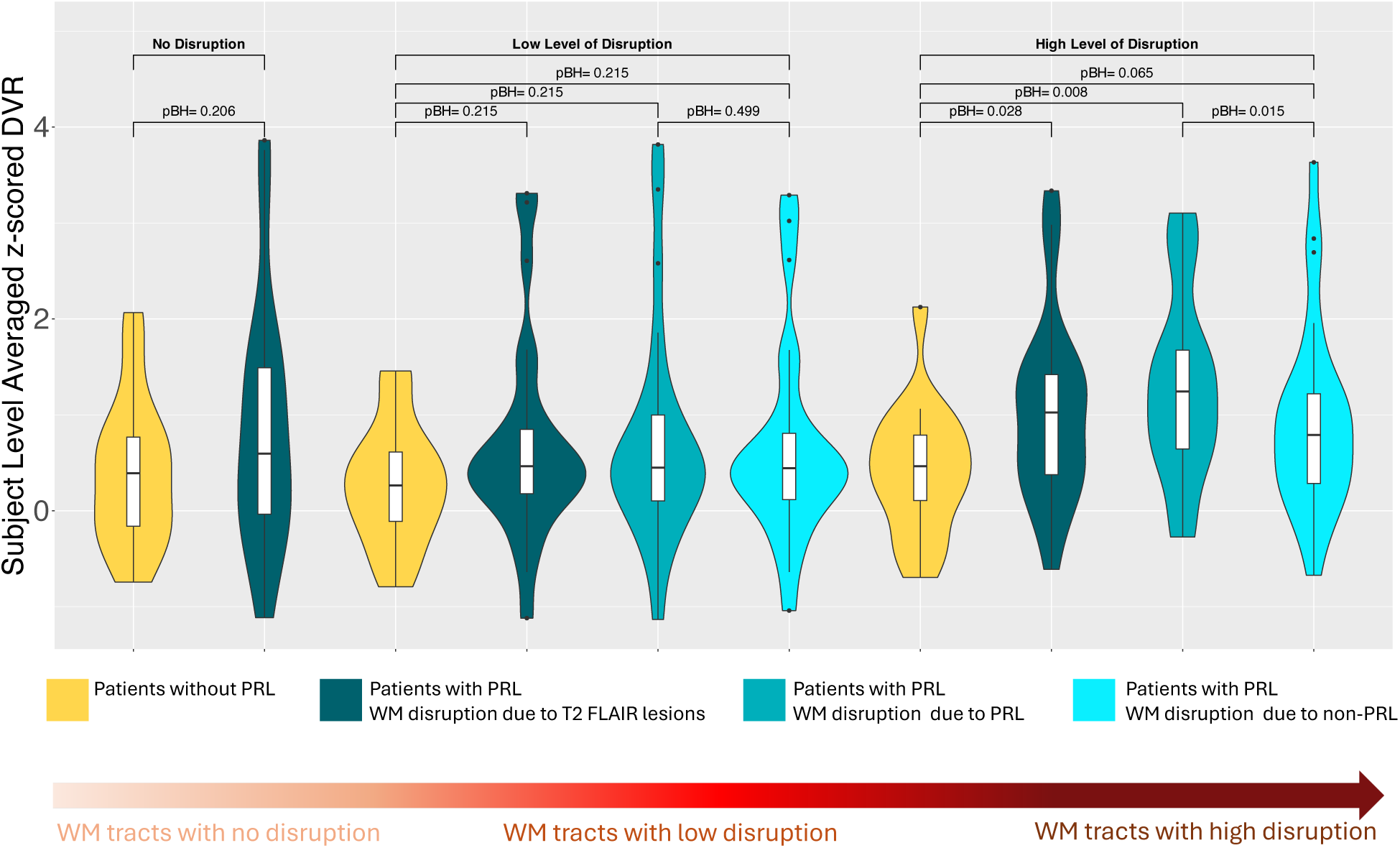
The patient-level z-scored DVR metrics (using age and sex-matched 11 HCs) averaged across the WM tracts where i) there was no disruption due to MS lesions (i.e. disruption in the WM tracts is zero), ii) there was a low level of disconnection due to lesions (i.e. disruption in the WM tracts is between 0 and 0.1), and iii) there was a high level of disconnection due to lesions (i.e. disruption in the WM tracts is between 0.9 and 1). The voxel-wise z-scored DVR was created in the WM using the voxel-wise DVR in 16 healthy controls. Then, the patient-level averaged z-scored metrics were calculated for each MS patient separately. The disruption metrics were separated as disruption due to PRL and disruption due to non-PRL for patients with at least one PRL. The yellow color represents the patients without PRL and the blue colors represent the patients with at least one PRL. The BH-adjusted p-values were presented in the figures.

When the age and sex-matched HCs were used to z-scored DVR metrics, we obtained similar results (See Supplementary Figure 6). A higher EDSS score was associated with increased z-scored DVR in the WM tracts with a high level of disruption due to MS lesions in patients with PRLs (adjusted p=0.0329), but not in patients without PRLs (adjusted p=0.194). When the association between EDSS and z-scored DVR was investigated in all MS patients, there was a significant association between EDSS and z-scored DVR in the WM tracts with high-level disruption in all MS patients (adjusted p=0.0147).

**Supplementary Figure 6:**
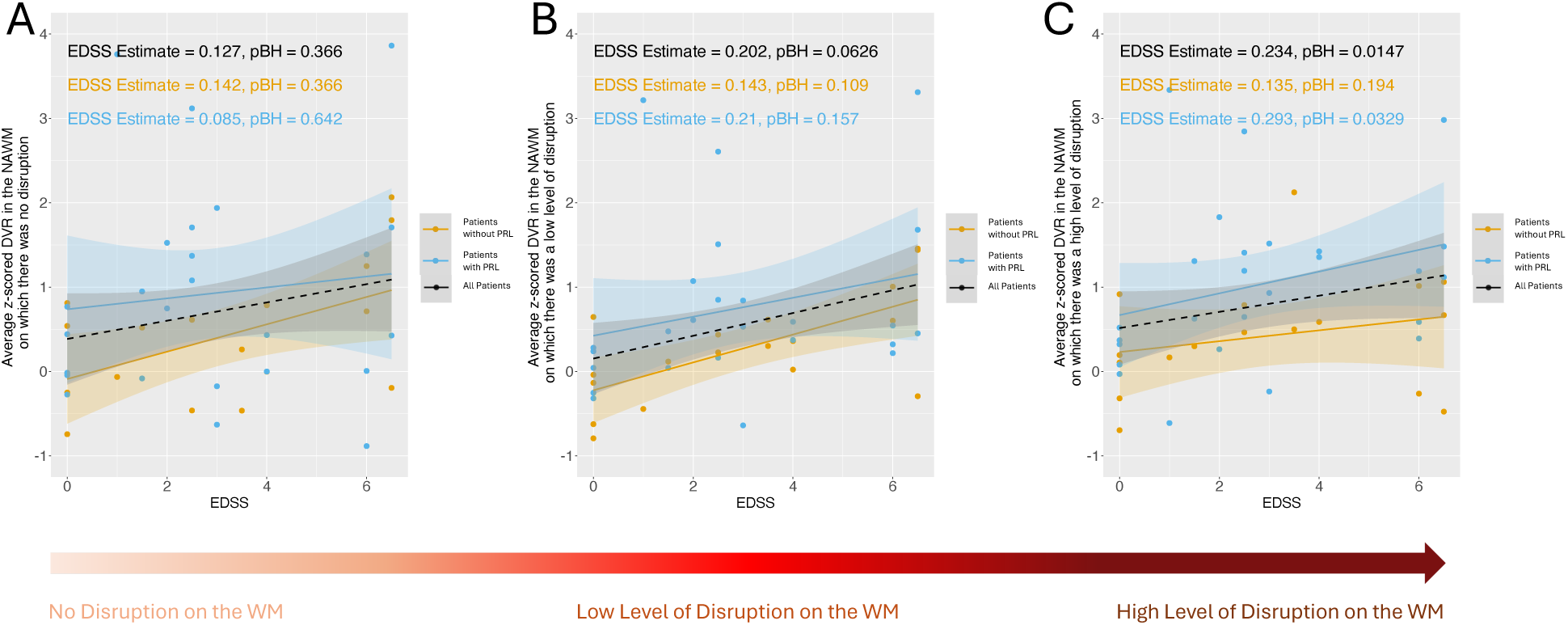
The scatterplots of EDSS and subject-level averaged z-scored DVR. The z-scored DVR was obtained using age and sex matched 11 HCs. We computed the subject-level averaged z-scored DVR in the WM tracts where A) there was no disruption due to MS lesions (i.e. disruption in the WM tracts is zero), B) there was a low level of disruption due to MS lesions (i.e. disruption in the WM tracts is between 0 and 0.1), and C) there was a high level of disruption due to MS lesions (i.e. disruption in the WM tracts is between 0.9 and 1). The estimates and the p-values associated with EDSS were calculated using a linear model where the output was the subject-level averaged z-scored DVR and the inputs were EDSS, age, sex, and MS type.

